# Alanine catabolism as a targetable vulnerability for MYC-driven liver cancer

**DOI:** 10.1101/2025.07.29.667471

**Authors:** Tonatiuh Montoya, Joyce V. Lee, Longhui Qiu, Abigail Krall, Nedas Matulionis, Yurim Seo, Brian N. Finck, Robin K. Kelley, Heather Christofk, Andrei Goga

## Abstract

Liver cancer is a leading cause of cancer-related death world-wide in part due to the shortage of effective therapies, and MYC overexpression defines an aggressive and especially difficult to treat subset of patients. Given MYC’s ability to reprogram cancer cell metabolism, and the liver’s role as a coordinator of systemic metabolism, we hypothesized that MYC induces metabolic dependencies that could be targeted to attenuate liver tumor growth. We discovered that MYC-driven liver cancers catabolize alanine in a GPT2-dependent manner to sustain their growth. GPT2 is the predominant alanine-catabolizing enzyme expressed in MYC-driven liver tumors and genetic ablation of GPT2 limited MYC-driven liver tumorigenesis. *In vivo* isotope tracing studies uncovered a role for alanine as a substrate for a repertoire of pathways including the tricarboxylic acid cycle, nucleotide production, and amino acid synthesis. Treating transgenic MYC-driven liver tumor mouse models with L-Cycloserine, a compound that inhibits GPT2, was sufficient to diminish the frequency of mouse tumor formation and attenuate growth of established human liver tumors. Thus, we identify a new targetable metabolic dependency that MYC-driven liver tumors usurp to ensure their survival.

## Introduction

Liver cancer is a leading cause of cancer-related death, with over 750,000 mortalities annually worldwide (1). Combination immunotherapy regimens (2–4) are the main treatments for surgically unresectable primary liver tumors. However, while a small subset of patients can experience deep and durable responses with combination immunotherapy, the majority of patients progress and succumb to the disease, thus motivating our search for new targeted therapies for this tumor type. Activation of MYC-dependent signaling is a pervasive event in liver cancer and more than 30% of hepatocellular carcinoma cases exhibit focal amplification or copy-number-gain of the MYC locus (5). Since clinically relevant MYC inhibitors are lacking, we and others have sought to identify druggable pathways selectively essential for the survival of MYC-overexpressing cells (6, 7). As a transcription factor, MYC can reprogram liver tumor metabolism by activating pathways such as glutaminolysis and glycolysis to meet the bioenergetic and biosynthetic requirements of cancer cells (8). We therefore hypothesized that identifying additional metabolic pathways selectively essential for MYC-dependent cancer cell growth (i.e. synthetic-lethal with MYC overexpression) could uncover new therapeutic targets.

Alanine is the second most abundant amino acid in circulation, second only to glutamine, and can be used by the liver to generate glucose via gluconeogenesis, which is then secreted to fuel muscle contraction in a process termed the glucose-alanine cycle (9). Alterations in alanine metabolism are also implicated in diseased states; for instance hepatic glutamic-pyruvic transaminase 2 (GPT2) levels increase in diabetes leading to augmented gluconeogenesis and hyperglycemia (10, 11). Key studies have also uncovered a role for tumor microenvironment-derived alanine in driving Kras-driven pancreatic cancer and have described a role for alanine transporters in directing the flow of alanine from pancreatic stellate cells to tumor cells (12, 13). However, the *in vivo* dependence on GPT2 and alanine metabolism in cancers from other tissue origins or with other oncogenic drivers is not known. Whether MYC-driven liver cancers are reliant on alanine and if this pathway can be targeted with pharmacological inhibitors could reveal potential new therapeutic approaches.

Here, we describe a role for alanine catabolism in sustaining MYC-driven liver tumors. We found that MYC-driven liver cancer cells use alanine to promote their growth in a GPT2-dependent manner both *ex vivo* and in a mouse model of MYC-driver liver cancer. We uncovered that alanine was a substrate for numerous metabolic pathways important for cancer cell growth, including the tricarboxylic acid cycle (TCA) cycle, non-essential amino acid synthesis, and nucleotide production. Additionally, we found that L-Cycloserine, an orally bioavailable compound that inhibits GPT2 (14, 15), slowed liver cancer cell growth *ex vivo* and *in vivo* in both transgenic mouse models and human liver cancer xenografts. Thus, GPT2-dependent alanine catabolism is exploited by MYC-driven liver tumors to enable their survival during nutrient deprivation, and inhibiting this pathway with L-Cycloserine may be efficacious in patients with liver cancer.

## Methods

### Conditional transgenic model of MYC-driven liver cancer

The LAP-tTA; TetO-cMYC; FVBN (LT2-MYC) mouse model has been previously described (16). LT2-MYC breeders were maintained on doxy chow to repress oncogene expression (200 mg/kg, Bio-Serv, ad libitum feeding) and taken off doxy chow at weaning to turn on MYC expression and induce tumor formation. Tumors typically form within 5-10 weeks, and ethical endpoint was determined by body score and abdominal palpation. Tumor samples were collected in OCT (Fisher Healthcare, 4585), 4% PFA in DPBS (Electron Microscopy Services, 15710), or snap frozen in liquid nitrogen.

For the L-Cycloserine tumor initiation study, LT2-MYC mice were taken off doxy chow at weaning and put on normal water or water with 500 mg/L L-Cycloserine at the same time (MedChemExpress, HY-B1122). Mice were monitored for tumor formation and sacrificed at ethical endpoint.

### Hydrodynamic model of MYC-driven liver cancer

The pT3-MYC-IRES-CRE vector was synthesized by GeneScript. The plasmid sequence was identical to that of the pT3-EF1a-MYC-IRES-LUC plasmid (Addgene, 129775), except the two LoxP sites were deleted, and the *Luciferase* coding sequence was replaced with the *Cre* coding sequence.

Hydrodynamic tail vein injection was done on wildtype male C57BL/6 (JAX, 000664) or GPT2 FLOX (10) mice of 6-8 weeks of age as done in reference (34). For every 2 mL of normal saline (Intermountain Life Sciences, Z1377), 13 µg of pT3-EF1-MYC-IRES-CRE, 13 µg px330-p53 (Addgene, 92046) and 520 ng of the pSB100X (Addgene, 34879) plasmids were dissolved and the solution then filtered through a.22 µm filter (Fisherbrand, 09-720-511). Mice were injected using 27.5 G butterfly needles (Terumo, SV*27EL) with a volume of plasmid solution equal to 10% of the mouse’s body weight in 5-8 seconds. Tumors formed within 5-10 weeks, and ethical endpoint was determined by body score and abdominal palpation. Tumor samples were collected in optimal cutting temperature (OCT), 4% PFA, or snap frozen.

### Xenograft model of liver cancer

HepG2 xenografts were established by injecting cells in Matrigel (Corning, 356234) into the hind flank of nude mice (JAX, 002019). HepG2 cells were lifted and resuspended at a concentration of 20 × 10^6^ cells / mL in DPBS (Gibco, 14190-144), and each mouse was injected with 100 µL of a 1:1 mix of Matrigel and cell suspension in DPBS using 25 5/8G syringes (BD, 309626). Tumor onset was monitored by caliper measurement of subcutaneous tumors. For tumor maintenance studies, once tumor diameter reached 1 cm mice were put on normal water or water with 500 mg/L L-Cycloserine which was relaced weekly. Mice were sacrificed once tumors reached 2 cm in diameter, and tumors were collected in OCT, 4% PFA, or snap frozen. Tumor volume was calculated using the formula V= (L*W^2^) / 2

### Histology and immunohistochemistry

Mice were sacrificed according to institutional protocols, then perfused with DPBS and tissues collected and fixed overnight in 4% paraformaldehyde (Electron Microscopy Sciences, #15700) on an orbital shaker. Tissues were then placed in 70% ethanol, and processed, stained, and imaged by Histowiz for c-MYC (Abcam, ab32072) and Hematoxylin and Eosin.

### Metabolomics

Steady state metabolomics of control livers and LT2-MYC tumors was performed and analyzed by Metabolon, as described in a prior study (16).

*In vivo* isotope tracing was performed on LT2-MYC mice that had been off DOX chow for 8-10 weeks. Tumor-bearing mice were anesthetized by isoflurane and kept on heating pads. Mice then had a catheter (Instech, C20PU-MJV2014) inserted into their jugular vein and tied in place using sutures (Surgical Specialties, SP117) and the catheter was then flushed with 50 μL heparin (McKesson, NDC 63739-931-14) to prevent coagulation. Mice were then infused with a ^13^C_3_ / ^15^N-alanine bolus of 0.114 mg per g body weight, followed by an infusion of 0.003 mg per g body weight per minute for 3 hours at a flow rate of 0.15 mL per hour using a Pump 11 Elite Dual Syringe Infusion Pump (Harvard apparatus, 70-4501). ^13^C_3_ / ^15^N-alanine was dissolved in sterile normal saline at 48 mg/mL. After infusion, mice were sacrificed, perfused with saline, and tumor and non-tumor tissue dissected and snap frozen.

Polar metabolite extraction from tissue samples was done by using a sharp blade to cut off a small piece of tissue 10-30 mg in weight. Each sample was placed in a 2 mL tube with a 5 mm stainless-steel bead (Qiagen, 69989) and 1 mL of methanol extraction solution composed of 80% methanol: 20% H_2_O with 100 nM trifluoromethanesulfonate (Fisher, A456-500; Fisher, W6-212; Sigma, 422843-5G). Samples were then extracted on a TissueLyser LT (Qiagen, 85600) by running samples for 1 minute at max speed, followed by a 1 minute break, three consecutive times. After extraction, samples were rested at −20°C for 10 minutes, and then vortexed and centrifuged at 17,000 x g for 10 minutes at 4°C. 700 µL of the supernatant was then transferred to new tubes and centrifuged in the same manner again. The top 500 µL of the supernatant was then transferred into a new tube and a volume of sample corresponding to a tissue equivalent of 4 mg was transferred into a new tube for evaporation. Samples were run on a DNA Vac-200 on high for 1-2 hours, or until the sample was completely evaporated. Samples were then stored at −80°C until they were run on the LC/MS.

Dried metabolites were reconstituted in 100 µL of a 50% acetonitrile (ACN) 50% dH20 solution. Samples were vortexed and spun down for 10 min at 17,000g. 70 µL of the supernatant was then transferred to HPLC glass vials. 10 µL of these metabolite solutions were injected per analysis. Samples were run on a Vanquish (Thermo Scientific) UHPLC system with mobile phase A (20mM ammonium carbonate, pH 9.7) and mobile phase B (100% ACN) at a flow rate of 150 µL/min on a SeQuant ZIC-pHILIC Polymeric column (2.1 × 150 mm 5 μm, EMD Millipore) at 35°C. Separation was achieved with a linear gradient from 20% A to 80% A in 20 min followed by a linear gradient from 80% A to 20% A from 20 min to 20.5 min. 20% A was then held from 20.5 min to 28 min. The UHPLC was coupled to a Q-Exactive (Thermo Scientific) mass analyzer running in polarity switching mode with spray-voltage=3.2kV, sheath-gas=40, aux-gas=15, sweep-gas=1, aux-gas-temp=350°C, and capillary-temp=275°C. For both polarities mass scan settings were kept at full-scan-range = (70-1000), ms1-resolution=70,000, max-injection-time=250ms, and AGC-target=1E6. MS2 data was also collected from the top three most abundant singly-charged ions in each scan with normalized-collision-energy=35. Each of the resulting “.RAW” files was then centroided and converted into two “.mzXML” files (one for positive scans and one for negative scans) using msconvert from ProteoWizard. These “.mzXML” files were imported into the MZmine 2 software package. Ion chromatograms were generated from MS1 spectra via the built-in Automated Data Analysis Pipeline (ADAP) chromatogram module and peaks were detected via the ADAP wavelets algorithm. Peaks were aligned across all samples via the Random sample consensus aligner module, gap-filled, and assigned identities using an exact mass MS1(+/-15ppm) and retention time RT (+/-0.5min) search of our in-house MS1-RT database. Peak boundaries and identifications were then further refined by manual curation. Peaks were quantified by area under the curve integration and exported as CSV files. If stable isotope tracing was used in the experiment, the peak areas were additionally processed via the R package AccuCor 2 to correct for natural isotope abundance. Peak areas for each sample were normalized by the measured area of the internal standard trifluoromethanesulfonate (present in the extraction buffer) and by the number of cells present in the extracted well.

### *In vitro* GPT activity assay

Tumors were homogenized in 2 mL tubes with stainless steel beads on a TissueLyser LT for 1 minute at max speed, followed by a 1 minute break, three consecutive times. For every mg of tissue, 10 µL of ALT assay buffer from the Alanine Transaminase Activity Assay Kit (Abcam, ab105134) was used to process the sample. Samples then were clarified by centrifugation at top speed at 4°C, and enzymatic activity assayed following the manufacturer’s instructions. For sample normalization, protein quantification was done using the DC protein assay kit (Biorad, 5000111).

### Cell culture

All cell lines were maintained in high glucose DMEM (Gibco, 11995-065) supplemented with 10% FBS (Gibco, 10437-028) and 100 µM non-essential amino acids (NEAA) (UCSF Cell Culture Facility) and were passaged every 2-3 days. EC4 cells were a gift from Dean Felsher (Stanford). HepG2 cells were purchased from ATCC. The PLC5, SNU475, and Tong cell lines were a gift from Xin Chen (UCSF).

All cell lines tested negative for mycoplasma, and human cell lines were validated by STR profiling (U Arizona Genetics Core).

### siRNA transfection

Non-targeting, mouse, and human *Gpt2*-targeting siRNAs (Dharmacon, D-001810-01-20; L-055921-01-0010, L-004173-01-000) were dissolved in RNAse-free H2O (Invitrogen, AM9938) at 20 µM and stored at −80°C in single use aliquots. siRNA knockdown was done by seeding cells at 100,000 (EC4 and Tong) or 150,000 (HepG2) per 6 well in 2 mL high glucose DMEM + 10%FBS + NEAA and transfecting 30 pmol of siRNAs per well with 5.5 µL of lipofectamine RNAi MAX (Invitrogen, AM9938) the following day, according to the manufacturer’s instructions. 24 hours post-transfection cells were lifted, passed through 70 µm cell strainers (Corning, 352350), and seeded for growth assays and knockdown validation. Importantly, prior to transfection, HepG2 cells were maintained at 70-90% confluency to ensure viability and growth post-transfection.

### Cell proliferation assays

For assays done on MYC high / low EC4 cells, we pretreated cells +/-10 ng/mL doxycycline (Fisher, BP2653-5) in high glucose DMEM + 10% FBS + NEAA for 3 days, changing half the media every 48 hours. For alanine growth assays done on the EC4, SNU475, PLC5, and TONG cell lines (Figures 1H-I, 1K), cells were cultured in DMEM no glucose / glutamine / pyruvate / phenol red (Gibco, A14430-01) supplemented with 10% dialyzed FBS (Gibco, 26400044), 5 mM or 500 µM glucose (Sigma, G8769-100ML), 500 µM glutamine (Sigma, G3126-100G), and 0 or 500 µM alanine (A7627-100G). Cells were seeded in clear bottom 96 well plates (Corning, 3603) in 200 µL media at a density of 1000 cells per well, and brightfield images were taken an hour after seeding to measure day 0 cell counts. Half the media was changed every other day, cells were then stained 3 days after seeding with PI (Invitrogen, P3566) and Hoechst 33342 (Biotium, 40046) at 1 µg/mL and 100 ng/mL, respectively, and were imaged 4 days after seeding for Hoechst and PI fluorescence. Glutamine was dissolved at 200 mM in water and frozen in single use aliquots at −80°C.

**Figure 1:**
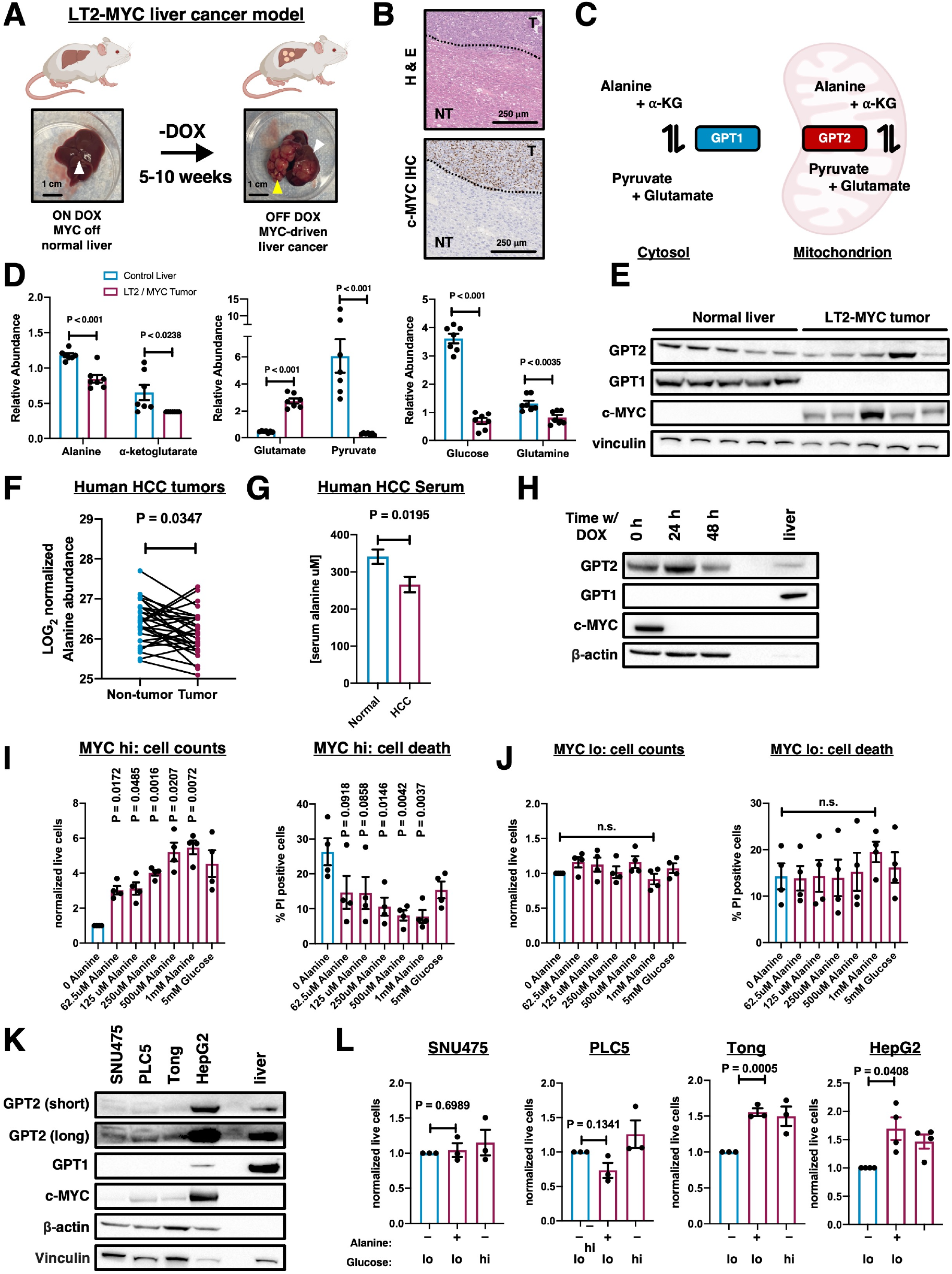
MYC-driven liver tumors exhibit a dependency on alanine metabolism. (A) Schematic of the MYC regulatable mouse model (LT2-MYC). Representative images of LT2-MYC livers with and without tumors. White arrows indicate non-tumor liver tissue, and yellow arrows indicate tumors. (B) Tissue sections from an LT2-MYC mouse were stained for hematoxylin and eosin or c-MYC and images taken at 5X magnification. Dotted lines indicate tumor boundary; “T” denotes tumor, “NT” denotes non-tumor. (C) Diagram showing compartmentalized activity of GPT1 in the cytosol and GPT2 in the mitochondrion. (D) LOG2 normalized abundance of glucose, glutamine, and alanine pathway metabolites by mass spectrometry in control liver versus LT2-MYC tumors. (n=7 for each group) (E) Immunoblotting of normal liver and LT2-MYC tumors for MYC and alanine enzymes. (F) LOG2 normalized abundance of alanine by mass spectrometry in tumor and adjacent nontumor tissue from human HCC (n=30 both groups), from Budhu et al. (G) Relative serum alanine abundance in healthy patients (n=10) and those with HCC (n=14), from Watanabe et al. (H) Immunoblotting of alanine enzymes and MYC in EC4 cells pretreated with doxycycline for 0-2 days to turn off MYC expression. Representative of 3 independent experiments. (I-J) Cell counts and cell death levels of MYC high (I) or MYC low (J) EC4 cells cultured in indicated alanine levels (n=4 independent experiments per condition). (K) Immunoblotting of MYC and alanine enzymes in a panel of human liver cancer lines. Representative of 3 independent experiments. (L) Proliferation response of a panel of human liver cancer lines to 500 μM alanine (n=3-4 independent experiments per line. Plots in (C), (G), (I), (J), and (K) show mean +/-SEM. Plot in (F) shows alanine abundance in matched tumor and non-tumor tissue from individual HCC cases. Significance in (D) and (G) determined using a two-sample t-test; matched pairs t-test in (F); multiple one-sample t-tests comparing alanine concentrations to control value of 1 with a Bonferroni correction in (I) and (J) for proliferation; one-way ANOVA with Dunnet’s multiple comparison test in (I) and (J) for cell death; one-sample t-test in comparing alanine to control value of 1 (L).

Alanine proliferation assays on HepG2 cells (Figure 1K) were done in 6 well plates (VWR, 10062-892) seeded at 28,000 cells per well. Cells were plated in 2 mL of DMEM no glucose / glutamine / pyruvate / phenol red supplemented with 10% dialyzed FBS, 5 mM or 500 µM glucose, 500 µM glutamine, 0 or 500 µM alanine, and half the media was changed every other day. After 4 days the supernatant containing non-adherent cells, and adherent cells were collected and counted by trypan blue staining (Invitrogen, T10282) on a countess cell counter. Glutamine was dissolved at 200 mM in water and frozen in single use aliquots at −80°C.

For alanine proliferation assays in Figures S1A-B, cells were cultured in DMEM no glucose / glutamine / pyruvate / phenol red supplemented with 10% dialyzed FBS, 5mM or 500 µM glucose, 0 or 500 µM alanine at the indicated glutamine levels. Tong cells were seeded at 1000 cells per well in clear bottom 96 well plates, and HepG2 cells were seeded at 5000 cells per well in Falcon 96 well plates (Corning, 353072) in 200 µL of the indicated media which was changed every 2 days. Tong cells were stained 3 days after seeding with PI (1 µg/mL) and Hoechst (100 ng/mL) and imaged 4 days after seeding for Hoechst and PI fluorescence. HepG2 cell counts were quantified by using the CYQUANT cell proliferation assay (ThermoFisher, C7026) 4 days after seeding. Glutamine was dissolved in water at 200 mM, stored at 4°C, and used within 2 weeks.

For siRNA proliferation assays in the EC4 cell line (Figure 3B), cells were collected 1 day post-transfection and seeded at 1000 cells per well in clear bottom 96 well plates in 200 µL DMEM no glucose / glutamine / pyruvate / phenol red supplemented with 10% dialyzed FBS, 500 µM glucose, 500 µM glutamine, 500 µM alanine, and brightfield images were taken an hour after seeding to measure day 0 cell counts. Methyl pyruvate (Thermo scientific, A13966.14) and dimethyl glutamate (Thermo scientific, L03764.06) were dissolved in water at 500 mM and added to the indicated conditions at 500 µM. Media was changed every 2 days, and cells were then stained 3 days after seeding with PI (Invitrogen, P3566) and Hoechst 33342 (Biotium, 40046) at 1 µg/mL and 100 ng/mL, respectively, and were imaged 4 days after seeding for Hoechst and PI fluorescence. siRNA proliferation assays in Tong cells (Figure 3D) were done identically, except cells were seeded in media containing 400 µM glutamine. siRNA proliferation assays in HepG2 cells (Figure 3D) were done similarly, except cells were cultured in media containing 350 µM glutamine and seeded in Falcon 96 well plates at 2500 cells per well. HepG2 cells were quantified by using the CYQUANT cell proliferation assay 4 days after seeding. For siRNA studies, glutamine was dissolved in water at 200 mM, stored at 4°C, and used within 2 weeks.

For knockdown validation, EC4, Tong, and HepG2 cells were seeded in high glucose DMEM + 10% FBS + NEAA at 50,000 cells per well in 6 well plates (EC4 and Tong) or 20,000 cells per well in 12 well plates (HepG2), and RNA was collected 2 days after seeding.

For L-Cycloserine dose response assays (Figure 4A), EC4 cells were pretreated +/-10ng/mL doxycycline in DMEM + 10% FBS + NEAA for 3 days, replenishing media every 2 days. For the proliferation assay, cells were seeded at 1000 cells per well in a clear bottom 96 well plate in 200 µL DMEM no glucose / glutamine / pyruvate / phenol supplemented with 10% dialyzed FBS, 500 µM glucose, 500 µM glutamine, and 500 µM alanine at the indicated L-Cycloserine levels. Brightfield images were taken an hour after seeding to measure day 0 cell counts. Media was changed every other day, and cells stained 3 days after seeding with PI and Hoechst at 1 µg/mL and 100 ng/mL, respectively. Cells were imaged on a Biotek high content microscopy plate reader for PI and Hoechst fluorescence 4 days after seeding. Glutamine was dissolved at 200 mM in water and frozen in single use aliquots at −80C.

To quantify cell proliferation in EC4, PLC5, SNU475, and Tong cells, PI and Hoechst 33342 puncta were counted, and total live cells were defined as day 4 (# Hoechst puncta - # PI puncta). We quantified cell growth as day 4 (# Hoechst puncta - # PI puncta) / day 0 brightfield counts. Cell death was quantified as (# PI puncta / # Hoechst puncta) x 100%. HepG2 cell proliferation in Figures S1B and 3D was quantified as day 4 CYQUANT cells counts / day 0 brightfield counts.

### High content microscopy

Cell growth assays were done on the Biotek Cytation 5 high content microscope (Agilent). The machine uses a Blackfly camera (BFLY-U3-2356M) equipped with a Sony IMX249 sensor. An Olympus 4X objective (UPLFLN4X) was used in conjunction with a Laser auto-focus filter cube (Biotek, 1225010), RFP filter cube (Biotek, 1225103) and a DAPI filter cube (Biotek, 1225100) to capture brightfield, PI, and Hoechst images.

### RNA extraction and cDNA synthesis

RNA was extracted using the QIAshredder Kit (Qiagen, 79656) and the Rneasy Mini Kit (Qiagen, 74106) according to the manufacturer’s instructions. cDNA was then synthesized using the High Capacity RNA-to-cDNA Kit (Applied Biosystems, 4387406).

### Quantitative PCR (qPCR)

qPCR reactions were set up using TaqMan Fast Advanced Master Mix (Applied Biosystems, 4444557) in optical 384 well plates (Applied Biosystems, 4309849). The following Taqman probes from Thermo fisher were used: Mm00558028_m1 Gpt2, Mm02619580_g1 Actb, Hs99999903_m1 ACTB, Hs00370287_m1 GPT2. Relative transcript levels were calculated using the ΔΔCt method.

### Immunoblotting

Proteins were extracted in RIPA buffer (50mM Tris-pH7.4, 150nM NaCl. 0.5% Triton, 0.2% Sodium deoxycholate) supplemented with protease and phosphatase inhibitors (Roche, 11697498001, 4906845001). Lysates were homogenized by passing them through 28.5 G insulin syringes (BD, 329424) 5-10 times and clarified by centrifuging at 17,000 x g for 15 minutes at 4°C. Supernatants were collected and transferred to new tubes, and protein content quantified using the DC assay (BioRad, 5000111). Samples were then prepared in 4X SDS sample buffer containing 100mM DTT (Fisher, BP172-25G).

Samples were run on 4–12% SDS-PAGE gels (Invitrogen, NW04122BOX) and transferred to nitrocellulose membranes (Invitrogen, IB23002). Membranes were blocked with 5% milk dissolved in Tris-Buffered Saline / 0.1% Tween 20 (TBST) for 1 hour at room temperature, and then incubated with primary antibodies dissolved in 5% milk / TBST overnight at 4*°* on a shaker. The following day membranes were washed 3 times in TBST for 5 minutes, and then incubated with HRP-conjugated secondary antibody diluted 1:5000 in 5% milk / TBST for 2 hr. After incubation, membranes were washed 3 times in TBST, and signal visualized using ECL (Bio-rad, 170-5061) or ECL Prime (Amersham, RPN2232).

The antibodies used and their dilutions in 5% milk / TBST follow: GPT1 (Abcam, ab202083, 1:1000), GPT2 (Proteintech, 16757-1-AP, 1:1000), GPT2 (Sigma, HPA051514, 1:1000), β-actin-HRP (Santa Cruz, sc-47778 HRP, 1:5000), MYC (Abcam, ab32072, 1:1000), Vinculin (Cell Signaling Technologies, 13901, 1:1000), α-Rabbit IgG-HRP (Promega, W4011, 1:5000).

## Results

### MYC-driven liver tumors exhibit a dependency on alanine metabolism

Cancers driven by the MYC oncogene exhibit tumor-specific metabolic phenotypes that can be inhibited to slow tumor growth, however additional vulnerabilities remain to be discovered. In order to find unexplored metabolic pathways that might be useful therapeutic targets for HCC, we used mass spectrometry to compare the global metabolite abundance of normal liver and MYC-driven liver tumors from the LAP-tTA; TRE-MYC (hereafter LT2-MYC) model, which has features of aggressive hepatocellular carcinoma (HCC) and hepatoblastomas (16). In this model, liver-specific MYC expression is induced when doxycycline is removed from the mouse diet, (i.e. OFF DOX) leading to tumor formation in 5-10 weeks (Figure 1A). Tumors formed in the LT2-MYC model are clearly demarcated from adjacent non-tumor on a macroscopic level (Figure 1A) and can distinguished by their cellular morphology and overexpression of MYC protein (Figure 1B). The glutamic-pyruvic transaminase isozymes GPT1 (cytosolic) and GPT2 (mitochondrial) catalyze the reversible transfer of an amino group from alanine to α-ketoglutarate, thus generating pyruvate and glutamate (Figure 1C). Notably, we observed that the abundance of the substrates of GPT1/2, alanine and α-ketoglutarate, were significantly reduced in LT2-MYC tumors as compared to normal liver, whereas glutamate, the product of GPT, accumulated significantly in LT2-MYC tumors (Figure 1D). Pyruvate, the other product of GPT, was not increased at steady-state in tumors, though the accumulation of pyruvate may be masked by its consumption through other pathways (e.g. lactate dehydrogenase, TCA cycle). Notably, glucose and glutamine, other metabolites used by MYC tumors to support their growth, were also depleted, suggesting that MYC tumors might depend on the alanine metabolic pathway.

We compared protein expression levels of GPT enzymes in normal liver and LT2-MYC tumors by immunoblotting and found that normal liver expressed both the cytosolic (GPT1) and mitochondrial (GPT2) isoforms of GPT, whereas LT2-MYC tumors mainly expressed mitochondrial GPT2 (Figure 1C, E). The observation that LT2-MYC tumors predominately express GPT2 suggests that they redistribute alanine metabolism to the mitochondrion, possibly to enable alanine usage by mitochondrial pathways such as the TCA cycle. Thus, based on the observation that LT2-MYC tumors have lower alanine levels than normal liver tissue, and specifically express mitochondrial GPT2, we hypothesized that alanine was being catabolized by liver tumors to promote their growth and survival.

The finding that alanine was depleted in a mouse model of liver cancer motivated us to investigate this pathway in human liver tumors. We therefore examined published liver cancer metabolomics datasets and observed that alanine abundance was consistently decreased in human liver tumors as compared to non-tumor liver across multiple studies (Figure 1F, refs. 17–19). Moreover, a prior study (20) examining amino acid levels in human liver cancer reported that alanine was depleted in the serum of patients with HCC as compared to healthy subjects (Figure 1G), possibly because of augmented liver tumor alanine catabolism and uptake from circulation. Collectively, the depletion of alanine in human serum, human liver tumors and in a transgenic MYC-driven liver tumor model led us to hypothesize that alanine could be a conserved metabolic substrate for liver cancer.

### Alanine promotes proliferation of MYC overexpressing liver cancer cells *ex vivo*

Having observed that alanine depletion is conserved in both a mouse model and in human liver cancer, we used EC4 cells (21), a tumor line derived from the LT2-MYC model, to interrogate the role of alanine metabolism *ex vivo*. Notably, when we compared alanine enzyme expression in EC4 cells treated with doxycycline to shut off MYC expression, we observed that mitochondrial GPT2 was expressed in EC4 cells, whereas cytosolic GPT1 was undetectable relative to normal liver lysates (Figure 1H), consistent with our observations in LT2-MYC tumors (Figure 1E). To test if alanine abundance is critical for the proliferation of EC4 cells, we cultured cells in nutrient levels that mimic the conditions of MYC liver tumors where glucose and glutamine are limited (Figure 1D) using DMEM + 10% dialyzed FBS, 500 µM glucose and 500 µM glutamine. The addition of alanine to MYC high EC4 cells resulted in a dose-dependent increase in viable cell counts and a dose-dependent decrease in cell death, as assayed by propidium iodide uptake (Figure 1I). In contrast, in EC4 cells that were pre-treated with doxycycline to turn off MYC expression, we did not observe appreciable changes in cell counts or cell death after 4 days of growth (Figure 1J) regardless of alanine supplementation. We therefore conclude that alanine promotes liver cancer cell proliferation and survival in a MYC-dependent manner, possibly through GPT2.

Based on the result that alanine could promote proliferation in a MYC-driven mouse liver cancer line, we next sought to generalize our findings to human cells. We used a panel of human liver cancer lines with varying levels of GPT1/2 and MYC expression which included the cell lines PLC5, SNU475, Tong, and HepG2. Notably, PLC5 and SNU475, which express lower levels of MYC and GPT2, did not exhibit a difference in cell counts after 4 days of growth in 500 µM alanine (Figure 1K-L). In contrast, the HepG2 cell line, which expressed the highest levels of MYC and GPT2, showed increased cell proliferation when growth media was supplemented with 500 µM alanine (Figure 1K-L). Notably, Tong cells, which expressed levels of GPT2 and MYC comparable to those of PLC5 cells, also showed increased cell proliferation when growth media was supplemented with 500 µM alanine (Figure 1K-L), suggesting that alanine might drive cell proliferation in other cellular contexts. We further sought to understand the dependency of human liver cancer cells on alanine by titrating levels of glutamine and measuring proliferation. We observed that alanine responsiveness depended on levels of glutamine present in cell culture, with HepG2 cells exhibiting the highest proliferative response when grown in 350 µM glutamine, and Tong cells in 400 µM glutamine (Figure S1A-B). We therefore concluded that liver cancer cells from both mice and humans can utilize alanine to increase proliferation *ex vivo*, partially compensating for limiting glucose and glutamine.

### Circulating alanine is metabolized by MYC-driven liver tumors *in vivo*

Our finding that MYC-expressing liver cancer cells use alanine to drive their proliferation and survival motivated us to understand the underlying mechanisms. We therefore established an *in vivo* isotope tracing strategy wherein tumor-bearing LT2-MYC mice were infused with a fully labelled ^13^C_3_ / ^15^N-alanine bolus and then continuously for 3 hours, similar to how other groups have traced metabolites in this tumor model (Figure 2A, ref. 22). Following infusion, tumor and non-tumor tissue were collected, polar metabolites extracted, and samples analyzed by mass spectrometry. We found that on average ~35% of the tumor alanine pool had at least one heavy isotope label (Figure 2B), thus confirming the efficacy of our tracing strategy. Notably, the ^13^C_3_ / ^15^N-alanine tracer made a greater contribution to the alanine pool in tumor as compared to non-tumor tissue, suggesting that MYC liver tumors might actively shuttle alanine to enable cancer cell catabolism. Having established and validated our *in vivo* tracing system, we next sought to understand the metabolic network downstream of alanine catabolism in MYC-driven liver tumors.

**Figure 2:**
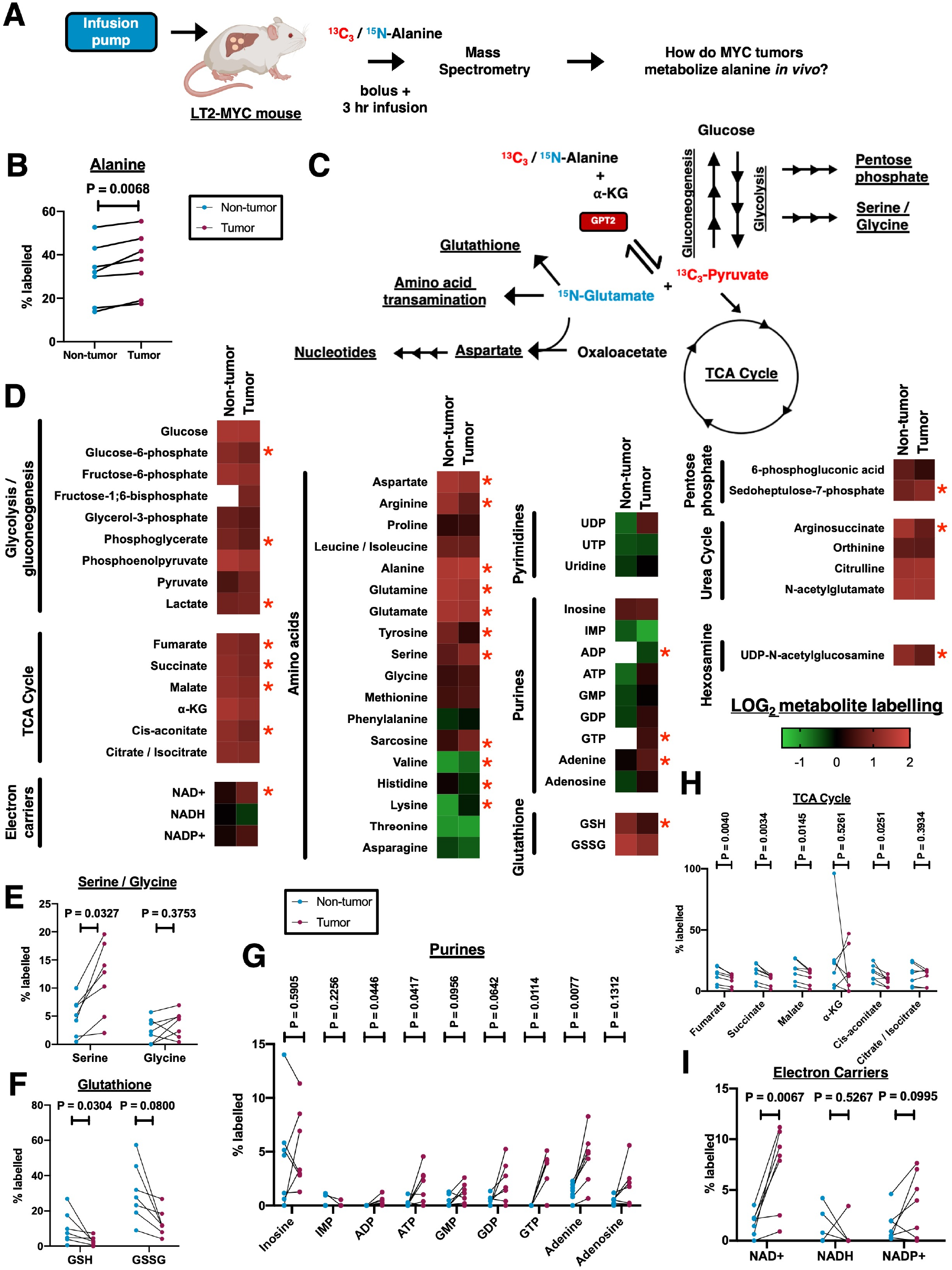
MYC-driven liver tumors exhibit a dependency on alanine metabolism. (A) Schematic showing the design of the *in vivo* isotope infusion study. (B) Contribution of ^13^C_3_ / ^15^N-alanine to the alanine pool in tumor and non-tumor tissue. (C) Diagram of the GPT enzymatic reaction and pathways that were labelled by the alanine tracer. ^13^C_3_ / ^15^N-alanine isotope labels are highlighted in red and blue, respectively. (D) Heat map showing LOG_2_ normalized average labelling efficiency of individual metabolites and their corresponding pathways. Red asterisks indicate P-value < 0.05. (E-H) Shown is the percentage of individual metabolites that had at least one label derived from ^13^C_3_ / ^15^N-alanine in pathways relating to serine / glycine synthesis (E), glutathione (F), purines (G), TCA cycle (H), and electron carrier molecules (I). n=7 mice for each condition. Significance in (B) and (D-H) was determined using a matched pairs t-test comparing labelling percentage in non-tumor and tumor tissue from the same animal.

We uncovered a repertoire of pathways downstream of alanine, with multiple metabolites being preferentially labelled in tumor versus non-tumor tissue. Among the pathways fed by alanine in tumors were metabolites relating to cellular bioenergetics, biosynthesis, and the oxidative stress response (Figure 2C-D). We observed that the labelling of serine was significantly higher in tumor than in non-tumor tissue (Figure 2E), consistent with prior work showing that serine synthesis is a MYC-stimulated process in hepatocellular carcinoma (23). We also observed labelling of glutathione (Figure 2F), consistent with our own work showing that the levels of the rate-limiting glutathione enzyme GCLC are indirectly repressed by MYC in liver tumors (16). Finally, we observed enhanced ^13^C labelling of lactate downstream of alanine (Figure S2A), consistent with the well-established role of MYC in promoting transcription of LDHA (24). Thus, our isotope tracing approach can identify known MYC-regulated metabolic pathways *in vivo*.

We next asked how tumor bioenergetics are altered in MYC tumors downstream of alanine. We observed that alanine-derived ^13^C labelled on average 10-20% of the pool of TCA cycle intermediates (Figure 2H), suggesting that alanine-derived ^13^C-pyruvate contributes to TCA cycle anaplerosis. Moreover, alanine labelling of the electron carriers NAD^+^ and NADP^+^ was higher in MYC tumors (Figure 2I), suggesting another mechanism by which alanine supports bioenergetics. Finally, we also observed that labelling efficiency of energy carriers ATP and GTP was higher in tumor versus non-tumor samples (Figure 2G). Thus, we find that alanine contributes to MYC-driven tumor bioenergetics in a multifaceted manner, both through driving TCA cycle anaplerosis, and the synthesis of bioenergetic molecules.

Given the role of alanine-derived pyruvate as a substrate for gluconeogenesis (9), we sought to understand how this pathway is altered in the context of alanine metabolism. We observed similar levels of ^13^C-pyruvate labelling by ^13^C_3_ / ^15^N-alanine in tumors and non-tumor tissue, suggesting that alanine supports pyruvate generation to similar extents in both contexts (Figure S2A). We then examined individual gluconeogenic intermediates and observed a reduction in ^13^C labelling of glucose-6-phosphate in tumors (Figure S2A), possibly due to increased conversion into intermediates of the pentose phosphate pathway, which is augmented in liver tumors (25). Additionally, we observed reduced ^13^C labelling of phosphoglycerate in tumor tissue (Figure S2A), possibly because of enhanced tumor serine synthesis, which uses phosphoglycerate as a substrate (23). Unexpectedly, we also observed that ^13^C-glucose labelling from alanine-derived pyruvate did not differ between tumor and non-tumors tissue, suggesting that MYC tumors retain a fully intact gluconeogenesis pathway. This is notable, given that the rate-limiting gluconeogenesis enzymes FBP and G6PC1 are generally downregulated in liver tumors (26–28), a trend which we also observed in the LT2-MYC tumor model (Figure S2B, ref. 29). This suggests that while gluconeogenesis is indeed repressed at the gene expression level, residual enzyme levels are still sufficient to sustain pathway activity in MYC tumors *in vivo*. Thus, we find that alanine supports gluconeogenesis in MYC tumors.

Given that pyruvate and glutamate, the products of GPT2, contribute to biosynthesis, we next asked if alanine differentially contributes to biosynthetic pathways in MYC tumors. We observed that alanine contributed to the production of most amino acids, consistent with the role of glutamate as an amino donor in transamination reactions (30). Specifically, we observed that serine and sarcosine had higher labelling in tumors, whereas aspartate, arginine, glutamine, glutamate, citrulline, and tyrosine had lower labelling in tumors (Figure S2D, H). Multiple products of purine synthesis were also labelled at a higher efficiency in MYC tumors, specifically ADP, ATP, GTP, and adenine (Figure 2G). Notably, multiple alanine-derived metabolites, including glutamine, glycine, and aspartate, as well as ATP, directly feed into the purine biosynthesis pathway, suggesting multiple mechanisms by which alanine supports purine synthesis (30). We also observed a contribution of alanine to the pentose phosphate pathway which generates ribulose-5-phosphate, a precursor of purine synthesis (31). Specifically, in tumors we observed higher labelling of the sedoheptulose-7-phosphate, and reduced labelling of glucose 6-phosphate (Figure S2G), the glycolytic intermediate from which the pentose phosphate pathway branches off, suggesting that alanine might coordinate pentose phosphate pathway activity and purine synthesis. Alanine also made a lesser contribution to the production of pyrimidines than to purines, with similar labeling efficiency (1-5%) of UDP, UTP, and uridine in tumor and non-tumor tissue (Figure S2E). Thus, our *in vivo* tracing studies support a model wherein MYC tumors catabolize alanine to sustain an interconnected network of pathways, ensuring necessary levels of energy production, biosynthesis, and glutathione production.

### MYC-driven liver tumorigenesis is dependent on GPT2 expression

We next sought to complement our descriptive isotope tracing studies with loss-of-function experiments to understand the mechanistic basis of alanine-dependent cell proliferation. Given that MYC-overexpressing cells have low levels of GPT1, we hypothesized that these cells are reliant on GPT2 for alanine-dependent cell growth in limiting glucose and glutamine conditions. To test this, we depleted GPT2 with pooled small interfering RNAs (siRNAs) or non-targeting siRNAs as a control (siNT) in EC4 cells. We measured total cell counts after culturing cells in limiting conditions of glucose and glutamine with 500 µM alanine supplementation for 4 days. We also supplemented cells with the cell permeable analogs of pyruvate (methyl-pyruvate) and glutamate (dimethyl-glutamate) at 500 µM to restore levels of the products of GPT2 and dissect their functional contribution to alanine-dependent cell proliferation downstream of GPT2 (Figure 3A). As expected, *Gpt2* transcript levels were reduced upon siRNA knockdown by quantitative PCR (qPCR), and we observed a significant reduction in total cell numbers in the vehicle-treated *Gpt2* knockdown condition as compared to siNT controls (Figure 3B-C). Moreover, methyl-pyruvate restored siGPT2 cell numbers to the levels of the siNT condition, and dimethyl-glutamate treatment also increased cell counts to a lesser extent (Figure 3B), suggesting that under these conditions cells depend more on GPT2-derived pyruvate than glutamate. We next sought to generalize our findings to the alanine-responsive human liver tumor cell lines Tong and HepG2. We observed similar reductions in total cell number upon *Gpt2* knockdown in HepG2 cells but did not observe a reduction in cell numbers in Tong cells (Figure 3D-E). Knockdown efficacy was also confirmed by qPCR in both lines (Figure 3D-E). Notably, Tong cells have much less MYC and GPT2 expression than HepG2 cells (Figure 1K); this suggests that alanine can promote the proliferation of human liver tumor cell lines via GPT2-dependent and independent mechanisms. Thus, our studies identify a dependence on GPT2 in alanine-driven cell growth in cells derived from the LT2-MYC model and in human liver cancer cells with high MYC expression.

**Figure 3:**
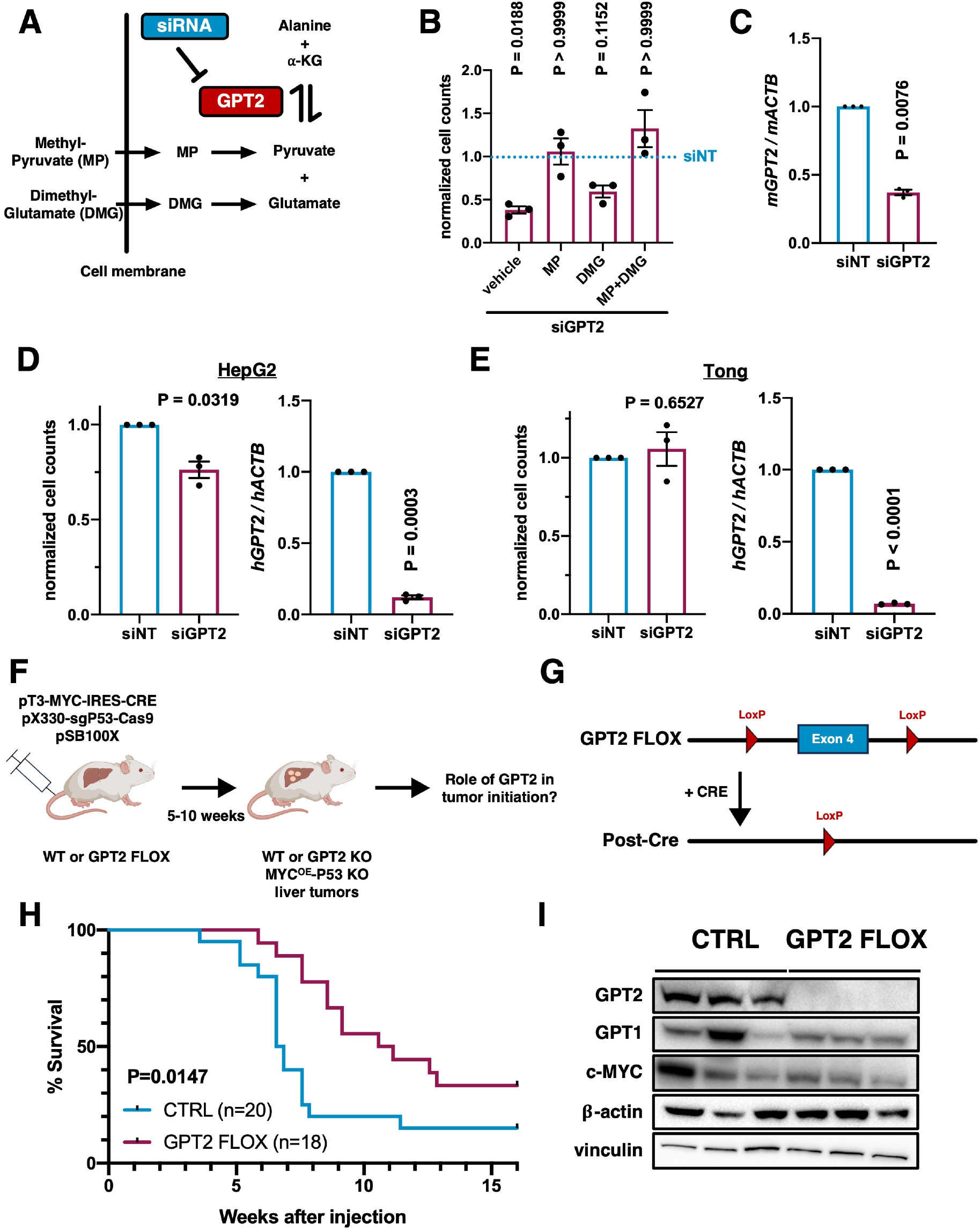
MYC-driven liver tumorigenesis is dependent on GPT2 expression. (A) Diagram of siGPT2 metabolite rescue study. (B) Proliferation response of EC4 cells +/-GPT2 knockdown in 500 µM alanine / glutamine media upon treatment with 500 µM methyl-pyruvate and / or dimethyl-glutamate (n=3 independent experiments each group). The blue dotted line represents the proliferation of control siNT cells which was normalized to 1. (C) Quantification of *Gpt2* knockdown efficiency in EC4 cells by qPCR (RNA from n=3 independent experiments). (D) Proliferation response of HepG2 cells to GPT2 knockdown in 500 µM alanine / 350 µM glutamine media, and qPCR knockdown validation of *Gpt2* knockdown (n=3 independent experiments). Proliferation and transcript levels of non-targeting control were both set to 1. (E) Proliferation response of Tong cells to GPT2 knockdown in 500 µM alanine / 400 µM glutamine media, and qPCR knockdown validation of *Gpt2* knockdown (n=3 independent experiments). Proliferation and transcript levels of non-targeting control were both set to 1. (F) Diagram of hydrodynamic tail vein transfection tumor initiation study. (G) Schematic showing how CRE deletes exon 4 of GPT2 leading to knockout of GPT2. (H) Survival of wildtype (n=20) versus GPT2 FLOX (n=18) mice upon liver tumor induction. (I) Expression profiling of wildtype and GPT2 knockout tumors. Plots in (B-E) show mean +/-SEM of normalized cell counts or normalized transcript levels where indicated. Significance in (B) determined multiple one-sample t-tests with a Bonferroni correction with the non-targeting control set to 1; a one-sample t-test with the non-targeting control set to 1 in (C-E); log-rank test in (H).

We next sought to test the requirement for GPT2 in the development of primary liver cancer. Towards this goal, we generated a transposon construct that allowed for the simultaneous expression of MYC and CRE recombinase, which we designated pT3-MYC-IRES-CRE. Either wildtype mice or mice which had a GPT2 allele in which the 4th exon was flanked by LoxP motifs (GPT2 FLOX, ref. 10) had tumor formation induced by somatic transduction of oncogenes via hydrodynamic tail vein injection. Mice received pT3-MYC-IRES-CRE, together with a construct to delete TP53 (pX330-Cas9-sgP53, ref. 32), and the sleeping beauty transposase (pSB100X, ref. 33) to generate tumors (Figures 3F-G, ref. 34). Thus, these tumors are driven by both MYC overexpression and TP53 loss and harbor either intact or deleted GPT2. We observed a marked difference in tumor burden in mice in which GPT2 was deleted; mice with GPT2 deletion also demonstrated a significant increase in median survival from 6.7 to 10.9 weeks compared to control GPT2 mice (Figure 3H). Notably, tumor penetrance also decreased from 85% to 66% in CTRL versus GPT2 FLOX mice (Figure 3H). We profiled protein expression in tumors generated in control and GPT2 FLOX mice and confirmed MYC overexpression and complete loss of GPT2 expression; we also failed to observe compensatory upregulation of GPT1 in GPT2 null tumors (Figure 3I). Thus, our studies support an *in vivo* functional role for GPT2 in MYC-driven liver tumor initiation.

### L-Cycloserine as a therapy for MYC-driven liver cancer

Our loss-of-function GPT2 studies indicated its potential as a therapeutic target in liver cancer and motivated us to investigate the utility of inhibiting GPT2 with the compound L-Cycloserine *in vivo*. While L-Cycloserine is not a specific inhibitor of GPT2 and may also inhibit the enzymes aspartate transaminase (AST) and serine-palmitoyl transferase (SPT) at higher concentrations (14, 15, 35), its oral bioavailability and ability to inhibit GPT2 motivated us to undertake preclinical studies to examine its potential as a therapy for MYC-driven liver cancer. To test this concept, we first used an *ex vivo* system wherein EC4 cells with low or high MYC expression were treated with increasing levels of L-Cycloserine. We observed that after 4 days of growth, EC4 cells with high levels of MYC exhibited a dose-dependent reduction in cell counts with increasing L-Cycloserine concentration whereas the proliferation of MYC low EC4 cells was not inhibited by increasing L-Cycloserine concentrations (Figure 4A). The MYC-dependence of L-Cycloserine sensitivity in cell-based models motivated us to investigate the *in vivo* efficacy of this compound in treating LT2-MYC tumors. MYC expression was induced in a cohort of LT2-MYC mice at weaning by removing doxycycline from their diet, and mice received water with 500 mg/L of L-Cycloserine to block GPT2 activity (Figure 4B). We used a fluorometric enzymatic assay to quantify GPT1/2 activity in the livers of mice treated with L-Cycloserine and confirmed that enzymatic activity was effectively abolished in drug-treated mice (Figure 4C). We then tested the efficacy of L-Cycloserine as a therapeutic by treating a cohort of mice in this manner for 8 weeks and sacrificed mice to assess tumor burden. We observed a pronounced difference in tumor burden as measured by liver mass, with control mice having an average liver weight over twice that of L-Cycloserine-treated animals (Figure 4D). Finally, we repeated a similar study but extended the duration to 3 months and found that L-Cycloserine improved time to ethical endpoint in drug-treated mice and markedly reduced tumor incidence from ~87% in CTRL to 30% in drug-treated mice (Figure 4E). Thus, our studies support the efficacy of L-Cycloserine treatment in reducing tumor burden in a mouse model of MYC-driven liver cancer initiation.

**Figure 4:**
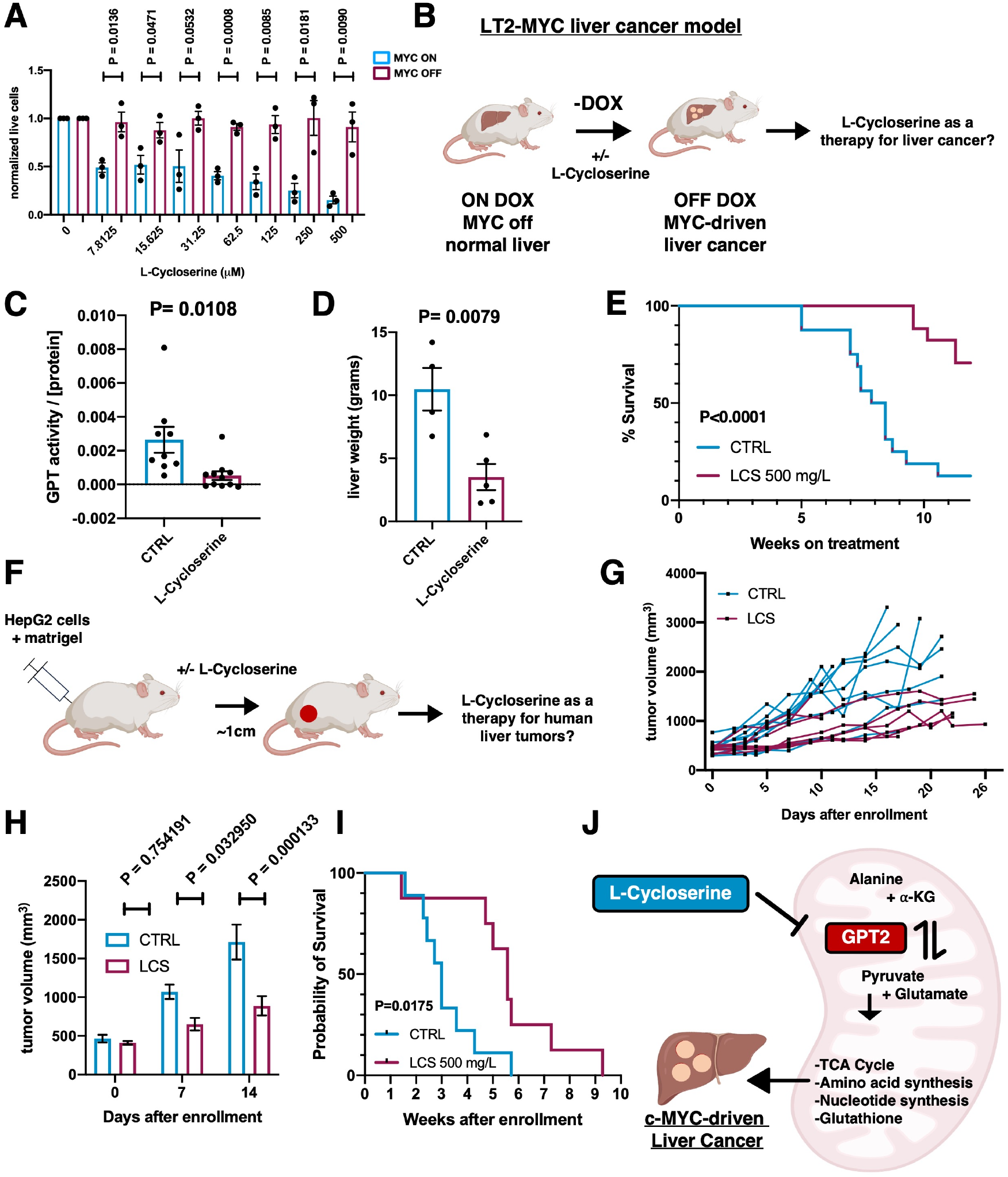
L-Cycloserine as a therapy for MYC-driven liver tumors. (A) Proliferation response of MYC high / low EC4 cells 500 μM alanine / glutamine media upon treatment with indicated L-Cycloserine levels as assayed by PI / Hoechst staining, with untreated control cells set to 1 (n=3 independent experiments each condition) (B) Diagram describing the L-Cycloserine tumor initiation study in the LT2-MYC model. (C) Quantification of in vitro GPT enzymatic activity in LT2-MYC tumors treated with 500 mg/L L-Cycloserine until ethical endpoint. (D) Tumor weights of LT2-MYC mice after 8 weeks of treatment with 500 mg/L L-Cycloserine in drinking water (n=4-5 mice each group). (E) Survival of LT2-MYC mice treated with or without 500 mg/L L-Cycloserine from the onset of MYC induction (n=16-17 mice per group). (F) Diagram describing the L-Cycloserine HepG2 tumor maintenance study. (G) Individual tumor volumes of HepG2 tumor xenografts after treatment with 500 mg/L L-Cycloserine or normal water control. (H) Average tumor volumes of HepG2 tumor xenografts after treatment with 500mg/L L-Cycloserine (n=7 mice) or normal water control (n=9 mice). (I) Survival comparison of HepG2 xenograft-bearing mice treated +/− L-Cycloserine. Endpoint was when tumors reached 2 cm in diameter. (J) Summary of the role of GPT2 in MYC driven liver tumorigenesis. Plots in (A), (C), (D), and (H) show mean +/− SEM. Significance in (A) and (H) determined using 2-sample t-tests with a Holm-Sidak multiple comparisons correction in (H); two-sample t-test in (C) and (D); log rank test in (E) and (I).

Based on the observation that L-Cycloserine was an effective therapy in a mouse model of liver cancer, we next sought to test the efficacy of L-Cycloserine in established human liver tumors. We generated mice bearing HepG2 xenografts, a model in which both MYC and GPT2 are highly expressed (Figure 1K), and treated mice with L-Cycloserine once tumors reached 1 cm in diameter (Figure 4F). We observed that the tumor volume of L-Cycloserine-treated mice was markedly diminished as compared to control tumors (Figure 4G-H) and that L-Cycloserine treatment extended the time for tumors to reach ethical endpoint (2 cm in maximal diameter; Figure 4I). Thus, these preclinical studies provide rationale for using L-Cycloserine, or other perhaps more selective inhibitors, as therapeutics in patients with MYC-driven liver tumors.

## Discussion

Liver cancer remains a major unmet public health need, and MYC overexpression demarcates an aggressive and difficult to treat subset of tumors (5, 36, 37). In this study we show that MYC-driven liver cancers are at steady state markedly depleted of the well-established metabolic substrates glucose and glutamine. Here, we find that MYC tumors usurp the machinery of the glucose-alanine cycle and catabolizes alanine in a GPT2-dependent manner to drive tumor formation. Our *in vivo* isotope tracing studies show that in addition to the role of alanine in driving gluconeogenesis, it can also drive bioenergetic and biosynthetic pathways required for MYC-driven tumorigenesis. Furthermore, we show that the compound L-Cycloserine, which inhibits GPT2, is efficacious at slowing the growth of established human liver tumors *in vivo* and preventing tumor formation driven by MYC. Thus, our study expands our understanding of how MYC reprograms tumor metabolism and identifies a new potential therapy for treatment of MYC-driven liver cancer.

Prior studies have described a role for alanine in Kras-driven pancreatic cancer, wherein stromal cells undergo autophagy and secrete alanine into the microenvironment, thus driving tumor growth. Our studies show that alanine catabolism can play a role in tissue and oncogene contexts beyond Kras-driven pancreatic cancer and suggest that other cancers arising in the liver or in the context of MYC overexpression may also catabolize alanine. Furthermore, our studies suggest that in addition to tumor microenvironmental alanine, the circulating systemic supply of alanine can also be leveraged as a metabolic substrate by liver tumors. Our data indicate that alanine catabolism is a druggable process and provide rationale for repurposing L-Cycloserine, or other clinically relevant GPT2 inhibitors, as a therapy for tumors that catabolize alanine.

These findings also suggest that MYC coordinates the generation and usage of multiple pools of pyruvate to ensure a balance between glycolysis and TCA cycle activity. MYC-dependent regulation of glucose transporters and glycolytic enzyme expression drives augmented glucose catabolism and lactate generation from pyruvate in the cytosol. This supports biosynthesis via shunting of glycolytic intermediates and regenerates NAD^+^ to maintain high rates of glycolysis (38, 39). In contrast, the generation of pyruvate via the mitochondrial-localized GPT2 enzyme (40, 41) might preferentially allow the usage of pyruvate by mitochondrial pathways such as the TCA cycle. Thus, MYC’s coordination of compartmentalized pyruvate generation could permit the simultaneous activity of the TCA cycle and glycolysis without the two processes having to compete for a limited supply of pyruvate.

It will also be important to understand how inhibition of alanine catabolism functionally interacts with other cancer therapeutics. For instance, T cells are alanine auxotrophs due to their lack of GPT enzyme expression and depend on extracellular alanine to sustain protein synthesis and T cell activation (42). As a result, inhibiting tumor cell alanine catabolism with L-Cycloserine could augment tumor microenvironmental levels of alanine, thus potentially promoting T cell activation and enhancing the efficacy of immune checkpoint blockade. Another open question is what the compensatory pathways are that enable tumor survival in the context of GPT2 inhibition. Pathways that generate the same metabolites as GPT2, for instance glutaminase (43) or other transaminases, could functionally compensate for GPT2 inhibition and may also be suitable targets for combinatorial metabolic therapy strategies.

Finally, while we observed efficacy of the compound L-Cycloserine *in vivo*, caution should be used in interpreting these results. L-Cycloserine is not a specific GPT2 inhibitor and inhibits aspartate transaminase and sphingosine-palmitoyl transferase at higher concentrations. Thus, the antitumor properties of L-Cycloserine may be due in part to inhibition also of other enzymes. Nevertheless, our studies demonstrate the utility of targeting GPT2 in liver tumors with both genetic and chemical approaches. The data presented here provide strong rationale for discovering more potent and specific GPT2 inhibitors which may prove to be useful as tools for research or as a therapy in the clinic.

Collectively, our work supports a model (Figure 4J) wherein GPT2-dependent alanine catabolism drives MYC-driven liver cancer by fueling tumor bioenergetics and supporting biomass generation. Our results also argue that inhibiting alanine catabolism with the compound L-Cycloserine, or other inhibitors of GPT2, could be an effective therapy for this aggressive and intransigent subset of liver cancer.

## Conflicts of Interest

The authors have no competing interests to declare.

## Data availability

Metabolomics data was uploaded to Metabolomics Workbench.

## Acknowledgements

We gratefully acknowledge support from the NSF Graduate Research Fellowship and a NIH NIGMS Initiative for Maximizing Student Development Fellowship (to T.M). Support was also received from Marcus Program in Precision Medicine Innovation (MPPMI), the Emerson Collective, Gazarian Family Foundation, and Bechtle Family (to A.G). P30 DK056341 (to B.F.) supported the generation of GPT2 floxed mice.

We thank H. Willenbring, L. Gilbert, and I. Jain for their mentorship and support of T.M. over the years. We also thank D. Van de Mark, R. Nakagawa, D. Superville, J. Gittins, F.C. Henry, and C. Pérez for helpful discussions. We also thank E. Atamaniuc for help with animal husbandry, and A. Chen for performing experiments.

Illustration of mouse with liver in Figure 1A created in BioRender. Goga, A. (2025). https://BioRender.com/mck8oko, and illustration of mouse in Figure 4F created in BioRender. Goga, A. (2025) https://BioRender.com/ccgm7z5. Illustration of liver and mitochondrion in Figure 4J also from BioRender.

## Author contributions

T.M. conceptualized the project, wrote the manuscript, designed, and contributed to experiments for all figures. J.V.L. contributed to experiments in Figures 2 and 4F-I, helped write the manuscript, and co-supervised the project. L.Q. performed microsurgery for the isotope tracing studies in Figure 2, and with S.S. performed hydrodynamic tail-vein injections for Figures 3F-H. A.K. and N.M performed mass spectrometry, data analysis, and helped in data interpretation for Figure 2. B.N.F. provided the gift of the GPT2 conditional knockout mouse and edited the manuscript. R.K.K. provided insight on human liver cancer and edited the manuscript. H.C. co-supervised the isotope tracing studies and provided feedback on the manuscript. A.G. supervised the project and helped write the manuscript.

---

**Supplementary Figure 1.**
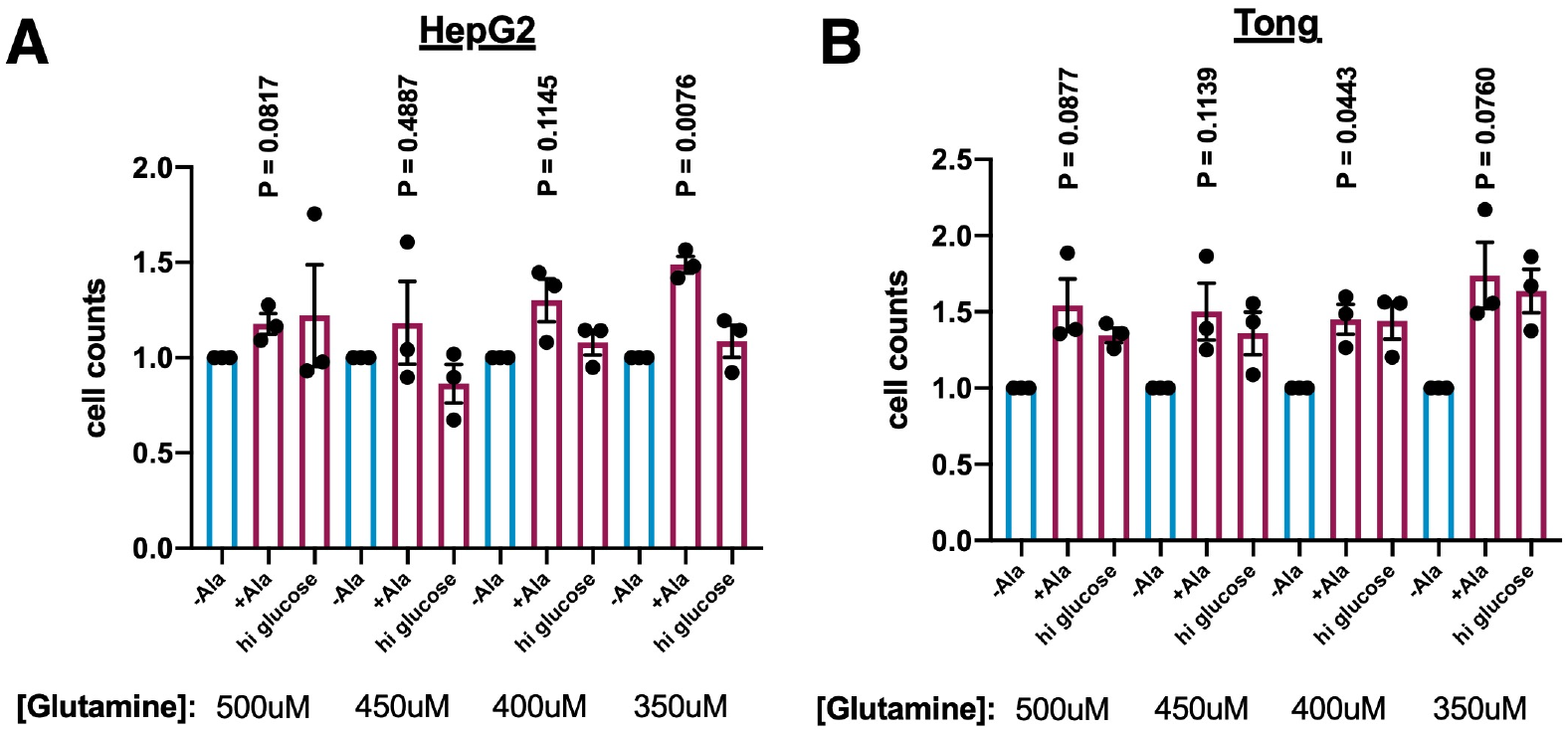
(A-B) Alanine proliferation response of HepG2 (A) and Tong (B) cells across a range of glutamine concentrations. All plots show mean +/-SEM with the 0 alanine condition for each glutamine condition set to 1. Significance in (S1A-B) calculated using multiple one-sample t-tests. n=3 independent experiments for each condition.

**Supplementary Figure 2.**
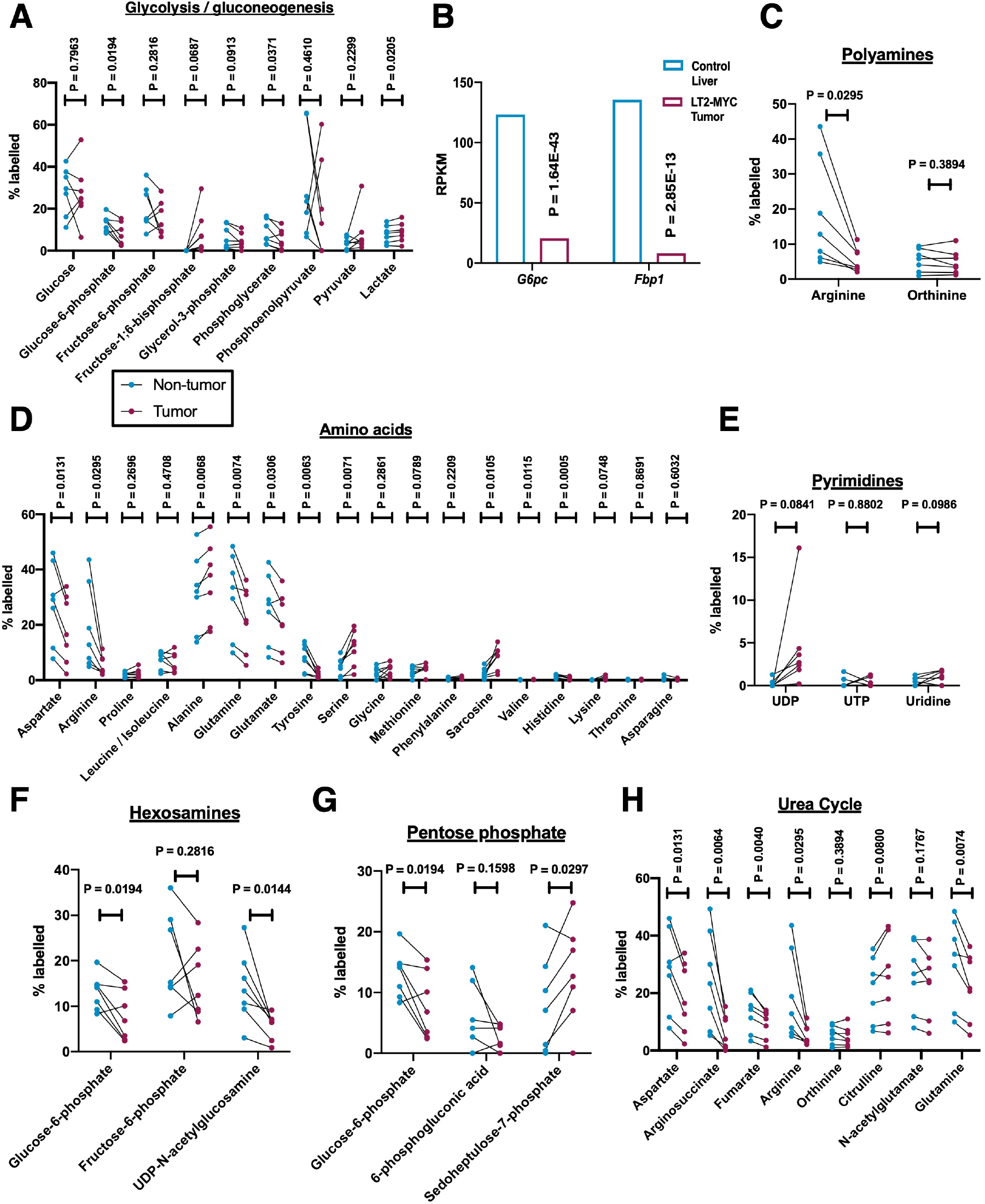
(A) Shown is the fractional contribution of ^13^C_3_ / ^15^N-alanine to glycolysis / gluconeogenesis (A), polyamines (C), amino acids (D), pyrimidines (E) hexosamines (F), pentose phosphate pathway (G), and the urea cycle (H). n=7 mice for each condition. (B) Transcript levels of *Fbp1* and *G6pc1* by RNA-seq, from ref. 40; n=11-16 per condition. Plots for S2A and S2C-H show metabolite abundance in matched tumor and non-tumor liver tissue for each animal. Plots in S2B show exonic RPKM values from RNA-seq. Significance for S2A, C-H was determined using a matched pairs t-test. Significance for S2B calculated in ref. 42, which used DESeq2 to calculate adjusted P-values for RNA-seq.

## Notes

### Competing Interest Statement

The authors have declared no competing interest.

### Summary of Updates

Small text changes and fixed one of the figure legends that had small errors.

